# Single-cell DNA methylation analysis tool Amethyst reveals distinct noncanonical methylation patterns in human glial cells

**DOI:** 10.1101/2024.08.13.607670

**Authors:** Lauren E. Rylaarsdam, Ruth V. Nichols, Brendan L. O’Connell, Stephen Coleman, Galip Gürkan Yardımcı, Andrew C. Adey

## Abstract

Single-cell sequencing technologies have revolutionized biomedical research by enabling deconvolution of cell type-specific properties in highly heterogeneous tissue. While robust tools have been developed to handle bioinformatic challenges posed by single-cell RNA and ATAC data, options for emergent modalities such as methylation are much more limited, impeding the utility of results. Here we present Amethyst, a comprehensive R package for atlas-scale single-cell methylation sequencing data analysis. Amethyst begins with base-level methylation calls and expedites batch integration, doublet detection, dimensionality reduction, clustering, cell type annotation, differentially methylated region calling, and interpretation of results, facilitating rapid data interaction in a local environment. We introduce the workflow using published single-cell methylation human peripheral blood mononuclear cell (PBMC) and human cortex data. We further leverage Amethyst on an atlas-scale brain dataset to describe a noncanonical methylation pattern in human astrocytes and oligodendrocytes, challenging the notion that this form of methylation is principally relevant to neurons in the brain. Tools such as Amethyst will increase accessibility to single-cell methylation data analysis, catalyzing research progress across diverse contexts.

## Main

Within an organism, thousands of distinct cell types are established and maintained using the same underlying genomic sequence. This diversity is largely mediated by the orchestration of gene expression through epigenetic modifications, including the covalent attachment of a methyl group to the 5’ position of cytosines by DNA methyltransferase proteins (DNMTs). Methylation can then facilitate the recruitment or inhibition of various transcription factors to bind DNA, thereby precisely tuning gene expression to the specific needs of each cell.^1^ Aberrant methylation patterns are associated with nearly every disease state - including many types of cancers, metabolic disorders, and neurological disorders^2–8^ – underscoring its critical relevance to a multitude of biomedical research fields.

Methylation canonically occurs at a cytosine followed by a guanine (mCG). In mammalian tissues, while the majority of available CG sites are methylated genome-wide, regulatory regions are CG-enriched and display highly variable methylation levels. It was more recently established that select cell types also exhibit substantial methylation at cytosines followed by non-guanine nucleotides (mCH).^9,10^ In mammals, this noncanonical pattern is most abundant in neurons, where mCH is rapidly deposited during peak synaptogenesis and can reach levels exceeding the number of mCG sites.^10,11^ The resulting patterns are more subtype-specific than mCG^11–13^ and are hypothesized to fine-tune expression of genes distinguishing closely related neuronal subtypes.^14^ While mCH is also present in glia at about five-fold lower levels,^9–12^ this population has historically been overlooked given the relative abundance in neurons, despite an initial report of NeuN^-^ cells exhibiting hyper-mCH in a gene subset with key roles in neuronal development.^11,12^

Subtype-specific methylation has also been challenging to study as single-cell methylation sequencing technologies have lagged behind other modalities like RNA or ATAC. While development of scBS-seq,^15^ scRRBS,^16^ and snmC-seq^17,18^ made it possible to analyze methylation on a single-cell level, caveats such as throughput, cost, and coverage still limited technology progression. These hurdles were recently ameliorated with the advent of combinatorial indexing-based strategies for methylation analysis (sciMET)^19–22^, an approach that exponentially increases throughput while simultaneously reducing costs per cell by orders of magnitude. The next challenge is data analysis: while comprehensive packages exist for exploration of single-cell RNA and ATAC data, available tools for single-cell methylation data analysis^23^ either 1) aren’t equipped to handle large-scale data; 2) are focused on mCG analysis only; or 3) address few steps in the scope of the entire workflow. One exception is ALLCools,^24^ a package written to analyze output of the snmC-seq^13,17,18^ workflow. However, ALLCools is Python-based, while a rich data analysis ecosystem exists within R framework due to the ubiquity of packages such as Seurat,^25–27^ Signac,^28^ Monocle,^29^ and ArchR.^30^

The objective of Amethyst is to provide a comprehensive R package for single-cell methylation analysis similar to what Seurat^25–27^ or ArchR^30^ have done for the RNA and ATAC fields. Amethyst is capable of efficiently processing data from hundreds of thousands of high-coverage cells in a relatively short time frame by performing initial computationally-intensive steps on a cluster followed by rapid local interaction of the output in RStudio. Versatile functions are provided to facilitate batch integration, doublet detection, clustering, annotation, differentially methylated region identification, and interpretation of results. Here we demonstrate the utility of Amethyst by exploring novel mCH patterns specific to mature glia of ectodermal lineage, a context previously emphasized in neuronal populations. Tools such as Amethyst will increase accessibility to single-cell methylation data analysis, catalyzing research progress.

## Results

### Amethyst enables resolution of biologically distinct populations

Following initial processing of reads, the first objective in single-cell methylation data analysis is to resolve distinct biological populations from gigabytes of base-level methylation calls (**Fig. 1a**). In Amethyst, this is done by first calculating methylation levels over a set of genomic regions for each cell followed by dimensionality reduction. Amethyst provides functions for a variety of genome windowing and scoring methods for this step depending on the tissue and method used. For example, when analyzing peripheral blood mononuclear cells (PBMCs), the mCG context is most relevant. Methylation patterns over fixed genomic windows, known regulatory regions, or established PBMC differentially methylated regions (DMRs) can all strongly resolve strong clusters (**Fig. 1b**). However, neurons uniquely have high levels of mCH, so clustering with information from both modalities provides better resolution than either context alone (**Fig. 1c**). The library preparation technique should also be taken into consideration for feature selection if results yield variable genome coverage; for example, the sciMET-cap^21^ method enriches for reads covering regulatory regions, so using targeted regions as features may resolve clusters better than fixed genomic windows. Amethyst and can also readily accept alternative strategies that can be implemented externally and imported into the object framework. This versatility will enable the rapid adoption of novel computational or molecular workflows as this fast-moving field evolves.

**Figure 1.**
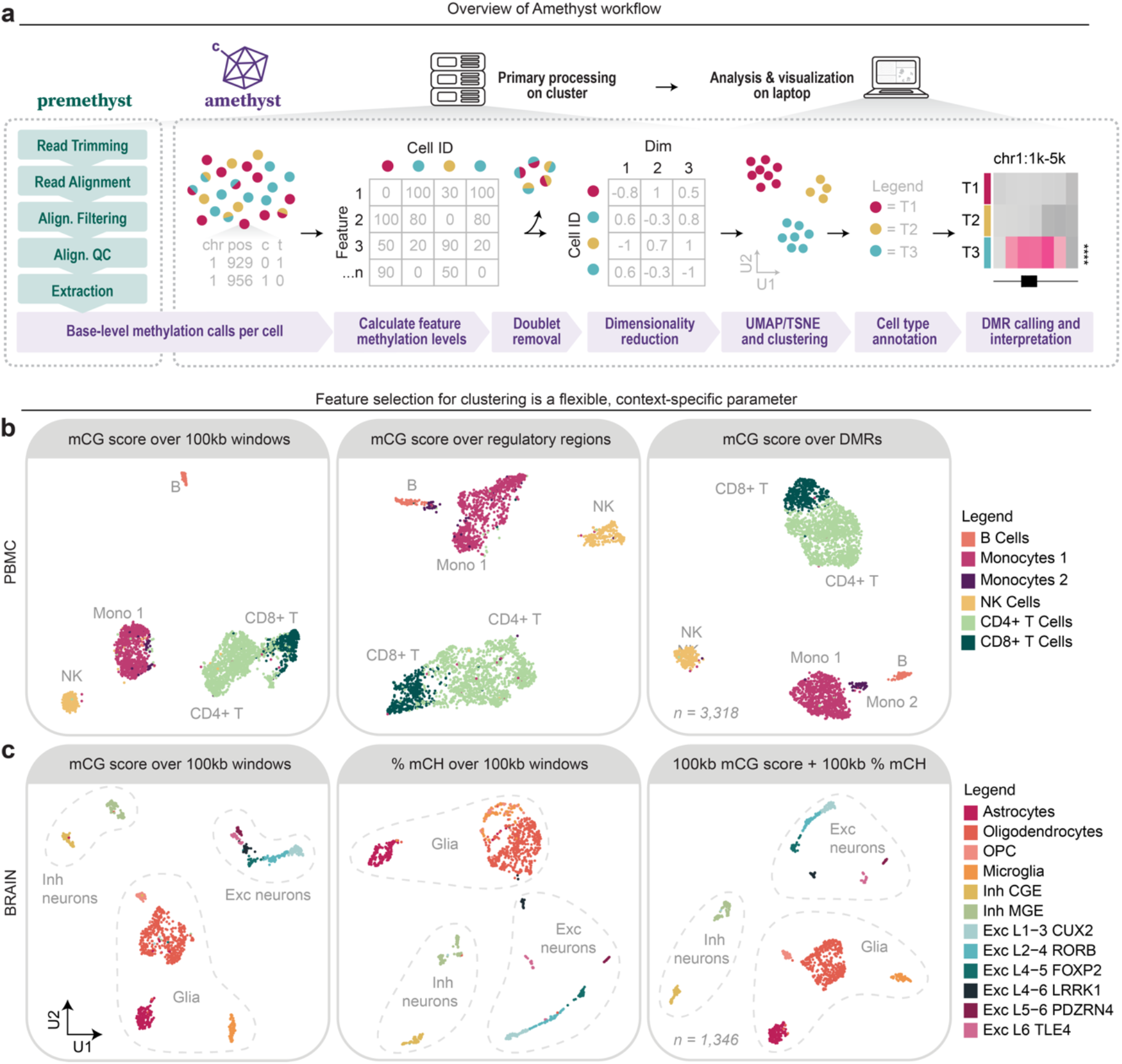
Introduction to Amethyst. **a**) Overview of the Amethyst workflow. The input format for Amethyst is generated from FASTQ files using processing pipelines such as Premethyst. Base-level methylation calls per cell are then used to calculate methylation levels over a feature set, which is used to resolve distinct biological populations. Following cell type annotation, differentially methylated regions (DMRs) can be determined and visualized. **b-c**) Examples of how using methylation levels over 100kb genomic windows, regulatory regions, or DMRs influences resolution of groups in human peripheral blood mononuclear cells (PBMCs) (**b**) and brain (**c**) data. In brain data, the mCH context is informative in addition to mCG.

Information contained in methylation levels over fixed genomic regions is then condensed to a lower dimensional space. Any dimensionality reduction can then be performed, but the default method for Amethyst is calculating fast truncated singular values with the implicitly restarted Lanczos bidiagonalization algorithm (IRLBA).^31^ Amethyst provides a helper function for estimating how many dimensions are needed to achieve the desired amount of variance explained. Batch correction with Harmony,^32^ mitigation of potential coverage biases, and doublet removal can be applied after this step if appropriate (**Fig. S1**). This output is then used to calculate cluster membership with the Louvain-based Rphenograph^33^ package and two-dimensional coordinates for visualization purposes with either uniform manifold approximation and projection (UMAP)^34,35^ or t-Distributed Stochastic Neighbor Embedding (t-SNE).^36^ Clustering results can be viewed and re-run in a local environment if populations are not effectively resolved.

**Figure S1.**
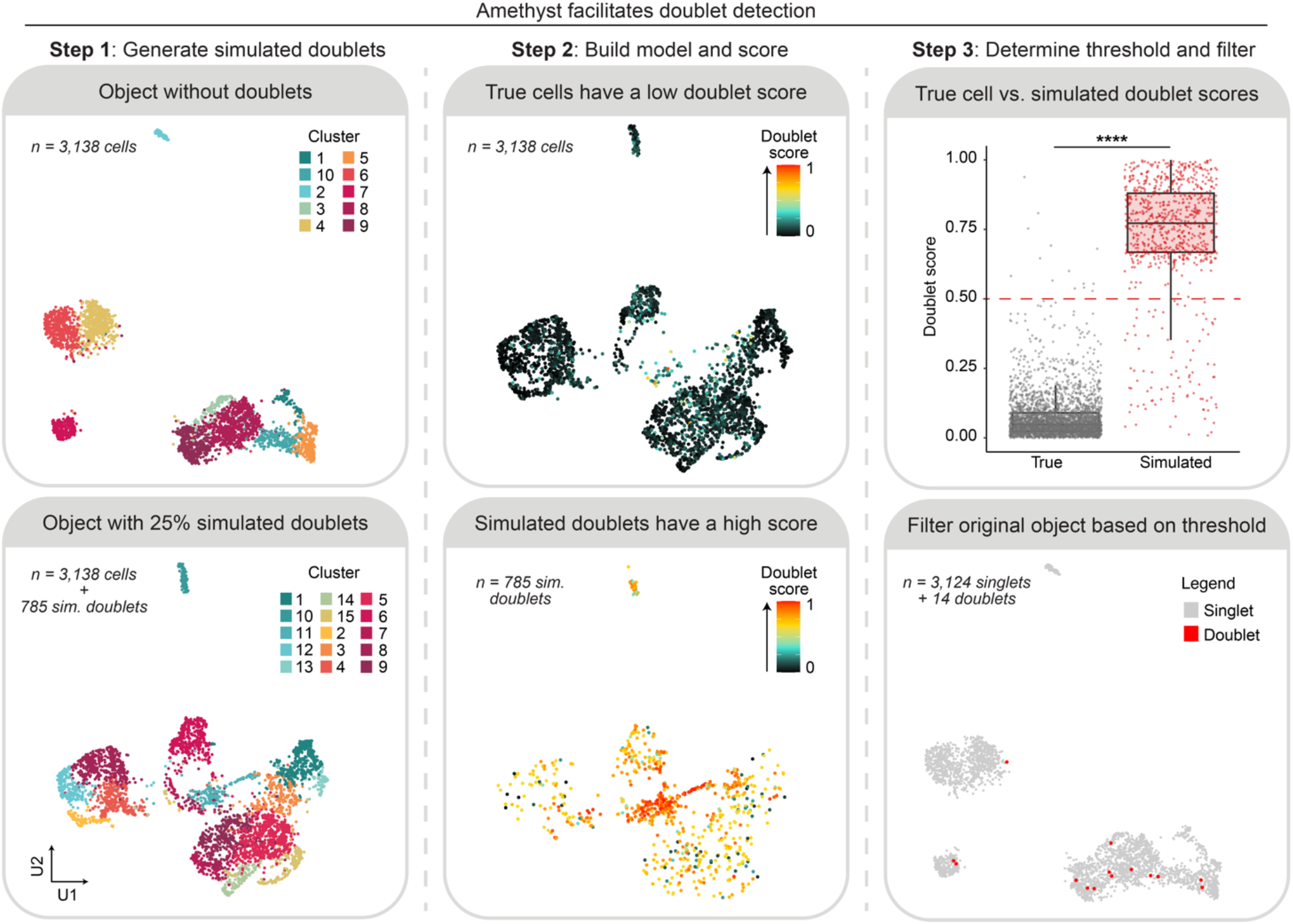
Amethyst enables removal of doublets. To identify doublets, a tunable proportion of simulated doublets are first constructed using an input matrix of methylation levels over a set of features (Step 1). A doublet model is then built with tools from the *caret*^37^ and *randomForest*^38^ packages and used to assign doublet scores (Step 2). In support of the method, artificial doublets have much higher doublet scores (mean±SE: 0.07± 0.001 vs 0.73±0.008; *W* = 2427350, *p* < 2.2E-16 using a two-sided Wilcoxon rank-sum test as *p*_Shapiro_ *<* 2.2E-16) and a higher fail rate (0.44% vs 12.95%; *X*^*2*^_1_ = 3284.8, *p* < 2.2E-16 using a two-sided Chi-square test) than true cells in this peripheral blood mononuclear cell example dataset. A threshold can be applied based on the doublet score distribution of true cells versus simulated doublets and used to remove predicted doublets from the original object (Step 3). Boxplot boundaries represent the first quartile, median, and third quartile values.

### Amethyst facilitates cell type identification through data interaction tools

Determining the biological identities of clusters can be challenging in any single-cell analysis, but particularly when exploring data from a novel modality. Amethyst provides multiple methods to facilitate this process including: 1) functions for visualizing mCG patterns across known marker genes (**Fig. 2a-b**); 2) tools to investigate gene body mCH or promoter mCG status between groups (**Fig. 2c-f**); and 3) comparison to existing references if available (**Fig. 2g**).

**Figure 2.**
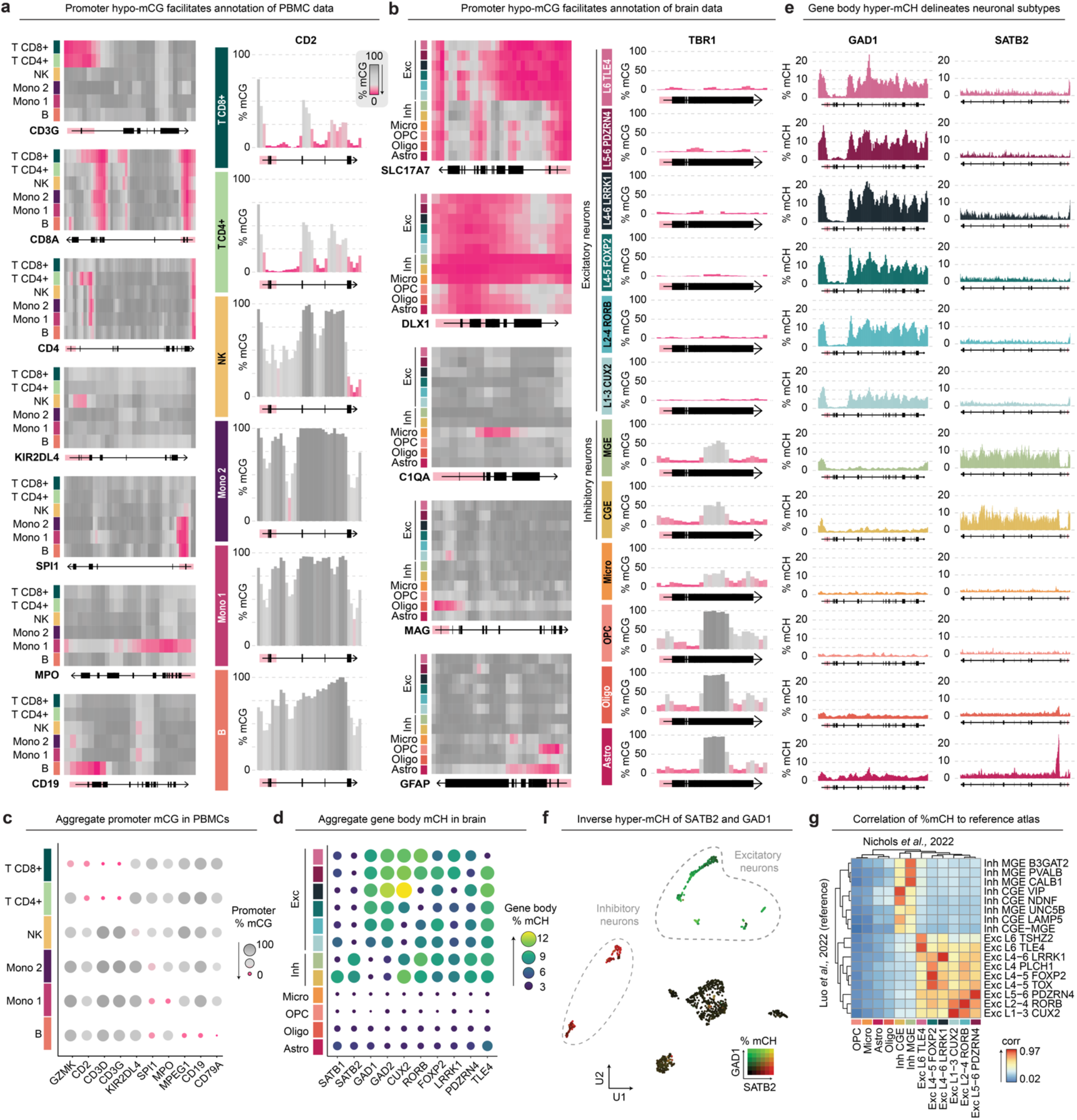
Amethyst facilitates cell type identification through data interaction tools. **a-b**) Visualization functions heatMap (left) and histograM (right) show mCG levels over canonical marker genes in human PBMCs (**a**) and brain tissue data (**b**). Color indicates %mCG for each 500bp genome window averaged by cell type with pink showing hypomethylated regions. Exons are in black and predicted promoter regions in red. **c**) The dotM function output shows promoter mCG levels of marker genes in PBMCs averaged by cell type. Both color and size indicate %mCG level, with pink being hypomethylated. **d**) The dotM function used to plot %mCH levels over the gene body in brain data averaged by cell type. Both color and size indicate gene body %mCH level, with yellow being hypermethylated. **e**) The histograM function used to show gene body mCH levels over canonical inhibitory marker *GAD1* and excitatory marker *SATB2*. Cell type headings in **b** apply. **f**) The dimM function shows a mutually exclusive mCH hypermethylation of *GAD1* (green) and *SATB2* (red) in UMAP space, which can be used to quickly classify inhibitory vs. excitatory neurons. **g**) Pearson correlation between average gene body mCH levels of 1502 relevant genes identified in Luo *et al*., 2022^12^ (rows) and data used in this study.^20^

Assessing mCG patterns across known marker genes is a powerful tool for cell type annotation. Hypo-mCG across the promoter region is canonically associated with gene expression, but this principle does not always apply: many validated marker genes will have no variable methylation patterns between groups; some will have universal promoter hypomethylation, yet a highly distinct pattern in the group expected to express the gene (for example, *SLC17A7* in **Fig. 2b**); and still others will appear to utilize a different promoter region than the bioinformatically predicted site. For all these reasons it is often more informative to visualize mCG patterns across the entire gene body. Amethyst provides two separate functions to do this - heatMap and histograM (**Fig. 2a-b**) – allowing the user to analyze patterns across genes in a far more nuanced manner than an aggregated metric. Consensus between known marker genes using this method is often sufficient for deduction of cell type annotation.

Amethyst also provides summary tools for calculating and visualizing mCG promoter levels if desired (**Fig. 2c**), but these are more powerful in the context of annotating brain tissue data, as neural populations arising from an ectodermal lineage can be annotated based on mCH alone (**Fig. 2d**). This distinct context accumulates postnatally over entire gene bodies and is anticorrelated with expression.^39^ mCH levels over canonical marker genes can easily be used to classify broad cell classes; for example, excitatory neurons have high mCH across inhibitory neuron enzyme *GAD1*, while inhibitory neurons have high mCH across excitatory neuron transcription factor *SATB2* (**Fig. 2d-f**).^11^ This context is highly subtype-specific,^11,40^ facilitating high-resolution annotation by methods such as anticorrelation to RNA datasets.^41^

Cell type annotation using gene-specific methylation information requires complex prior knowledge of expression patterns in the model system and puts significant weight on few genes. Users may wish to have a more automated cell type annotation process by comparing methylation over variable features or 100kb windows to an atlas, as the inclusion of thousands of features instead of a few canonical marker genes can produce clearer correlations. This method depends on the existence of comprehensive and carefully annotated datasets, which exist for brain data^11–13^ but few other tissues. We have utilized a landmark human brain atlas^12^ to demonstrate how correlation of mCH patterns over clusters facilitates cell type identity (**Fig. 2g**), but similar methods can be applied using the mCG context as well. Currently, Amethyst provides two built-in references to facilitate brain and PBMC data identification, and more tissue types will continue to be made available as the body of single-cell methylation atlases expands.

### Amethyst robustly identifies differentially methylated regions

For the majority of single-cell methylation experiments, successful identification of DMRs is at the core of analysis. Amethyst utilizes a smoothed sliding window approach of methylation levels over short genomic regions, overcoming sparsity challenges by aggregating observations of methylated and unmethylated cytosines per cluster. A variation of a Fisher’s exact test from methylKit^42^ is then used for each genomic window on the proportion of methylated vs unmethylated cytosines. Following strict correction for multiple testing, adjacent DMRs are collapsed and annotated with any overlapping genes. Amethyst functions such as heatMap can then be used to visualize DMRs within their genomic context (**Fig. 3a-c**).

**Figure 3.**
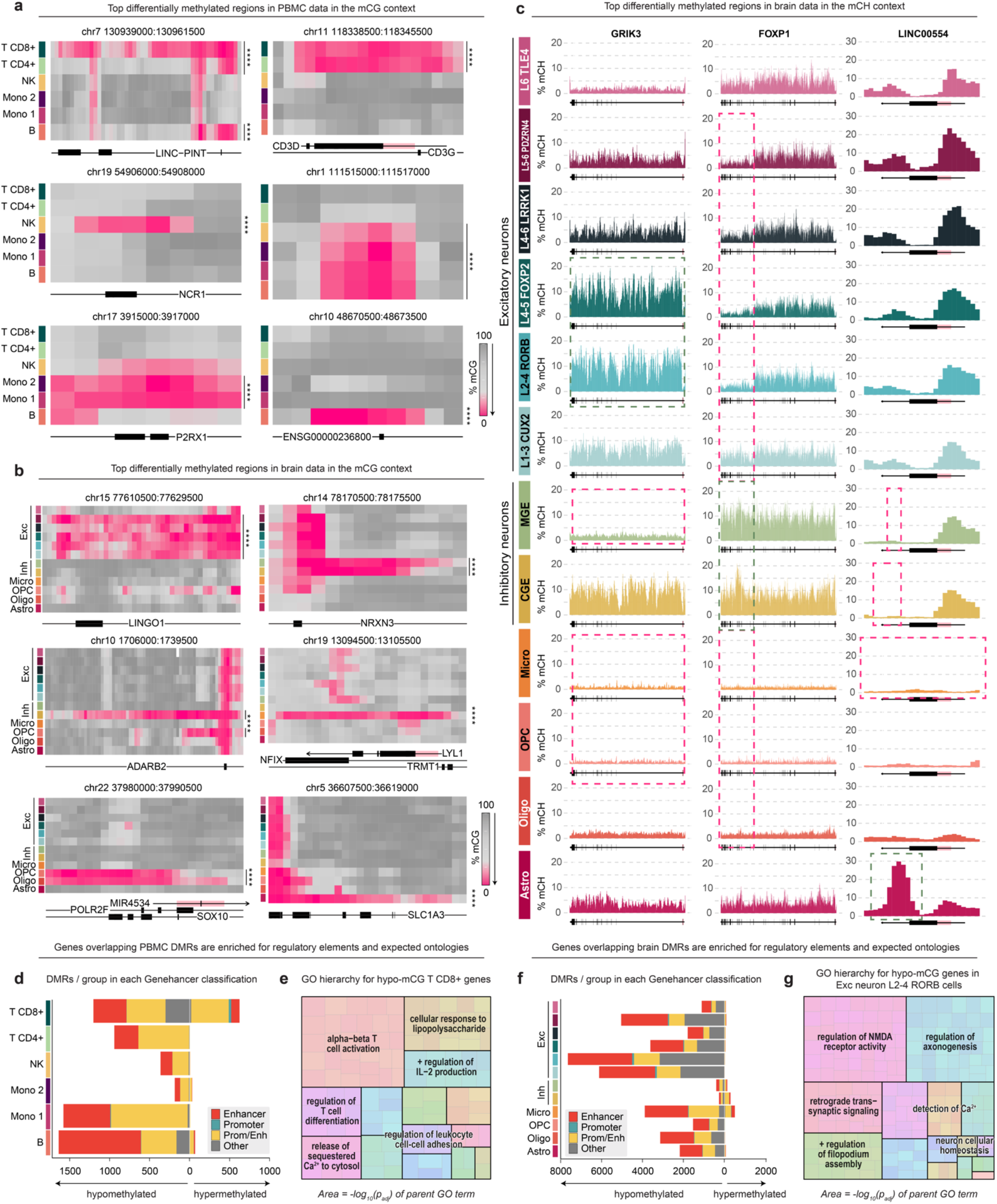
Amethyst robustly identifies differentially methylated regions (DMRs). **a-b**) heatMap of top hypomethylated DMRs identified in peripheral blood mononuclear cells (PBMCs, **a**) and brain tissue data (**b**). Color indicates %mCG for each 500bp genome window averaged by cell type with pink showing hypomethylated regions. Any overlapping genes are plotted below. Exons are in black and predicted promoter regions in red. Statistics were calculated using a variation of a two-sided Fisher’s exact test^42^ with Bonferroni adjustment for multiple testing (see **Methods**). **c**) histograM showing genes containing top DMRs identified in the mCH context for brain data. Key variable regions are highlighted with either a pink (hypomethylated) or green (hypermethylated) dashed box. The gene body is plotted below with exons in black and predicted promoter regions in red. Statistics were calculated using a variation of a two-sided Fisher’s exact test^42^ with Bonferroni adjustment for multiple testing (see **Methods**). **d**) GeneHancer^43^ annotation of PBMC DMRs divided by direction and cell class. **e**) Treemap showing hierarchical structure of gene ontology (GO) results for genes with hypomethylated regions identified in the CD8+ T cell group. Key parent terms are shown in black with nested related child terms inside the parent box. Area corresponds to -log_10_(*p*_elim_) as calculated with the *topGO*^44^ and *rrvgo*^45^ R packages using the “elim” method. **f**) Genehancer annotation of brain DMRs divided by direction and cell class. **g**) Treemap showing hierarchical structure of gene ontology (GO) results for genes with hypomethylated regions identified in the Exc neuron L2-4 RORB cell group. Key parent terms are shown in black with nested related child terms inside the parent box. Area corresponds to -log_10_(*p*_elim_) as calculated with the *topGO*^44^ and *rrvgo*^45^ R packages using the “elim” method.

In tests using published PBMC^21^ and brain^20^ datasets, top DMRs fall within canonical marker genes (**Fig. 3**), strongly supporting the efficacy of this method. For example, the top hypomethylated DMR in the CD4+ T cell group encompasses *CD3D* and *CD3G* (*p*_adj_ = 1.36E-122), which encode subunits of the T-cell receptor-CD3 complex and play essential roles in facilitating the adaptive immune response (**Fig. 3a**). In brain data, the top hypomethylated DMR for the astrocyte group falls across much of *SLC1A3* (*p*_adj_ = 1.22E-180), which encodes pan-astrocyte marker GLAST (**Fig. 3b**). This approach can directly be applied to the mCH context as well, revealing both previously known DMRs – such as glutamate ionotropic receptor subunit *GRIK3* and forkhead box transcription factor *FOXP1* – as well as unstudied avenues for exploration, such as a dramatic inverse hypermethylation pattern across long non-coding RNA *LINC00554* in astrocytes relative to neurons (significance across various groups and regions; see **Fig. 3c**).

To further confirm the validity of results, we assessed how many DMRs in the mCG context overlapped with regulatory elements, which are known to contain the highest variability in methylation levels. In PBMC data, 92.2% of DMRs fell within enhancer or promoter regions (**Fig. 3d**), despite only about 12% of the genome being annotated as such in GeneHancer.^43^ To test if DMRs were enriched for expected biological processes, we performed gene ontology (GO) analysis^46^ for genes with hypomethylated regions in the CD8+ T cell group. Result redundancy was simplified using the rrvgo^45^ package and plotted in a hierarchical treemap structure to obtain a more holistic view of results (**Fig. 3e**). Top parent terms were strongly related to T cell biology, including alpha-beta T cell activation (*p*_*elim*_ = 1.4E-04) and regulation of T cell differentiation (*p*_*elim*_ = 1.1E-04). Similar patterns were observed from DMRs identified in brain data: the majority (69.4%) fell within regulatory regions (**Fig. 3f**) and top ontology enrichments for genes with hypomethylated regions in the excitatory neuron L2-4 RORB cells were related to glutamatergic synaptic signaling processes. Examples include regulation of NMDA receptor activity (*p*_*elim*_ = 1.1E-04) and retrograde trans-synaptic signaling (*p*_*elim*_ = 1.3E-04; **Fig. 3g**). Altogether, the enrichment of DMRs for regulatory regions and expected GO terms supports the efficacy of Amethyst’s DMR identification approach.

### Differentially methylated gene analysis in Amethyst reveals a novel non-CG methylation signature in glia

The sliding-window DMR identification method is applicable to mCG analysis in any tissue, as mCG is variably deposited across regulatory elements in the genome. In neurons, mCH is known to accumulate abundantly across large TAD-guided domains and is thought to fine-tune gene expression by suppressing genes defining closely related neuronal subtypes.^11,14,39,47^ To demonstrate the utility of gene-specific mCH analysis, we used Amethyst to test all protein coding genes for cluster-specific variable mCH. In support of previous research,^10–12,48^ neuronal groups had orders of magnitude more hypermethylated genes than glia, and no hyper-mCH genes were identified in immature OPCs or mesoderm-originating microglia (**Fig. 4a**).

**Figure 4.**
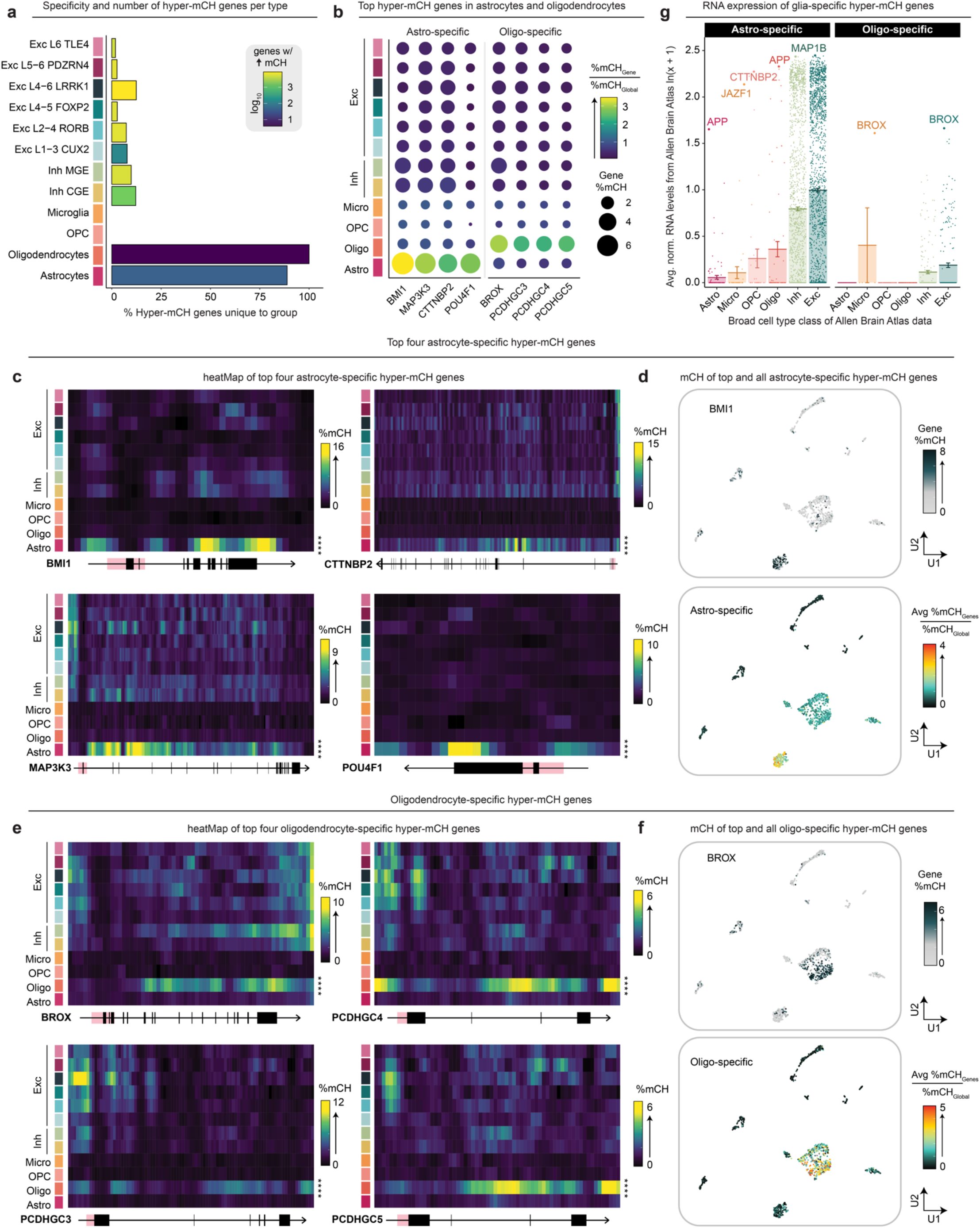
Differentially methylated gene analysis in Amethyst reveals a novel non-CG methylation signature in glia. **a**) Bar chart with color indicating the number of hypermethylated genes in the mCH context per group, and height showing the percentage of hits only found in that population. All protein-coding genes in the genome were tested for significance using a two-sided Wilcoxon rank-sum test with Bonferroni correction. **b**) Dot plot showing average mCH values of per cell type of the top four hyper-mCH genes found to be specific to the astrocyte and oligodendrocyte populations. Size indicates %mCH across the gene body; color shows this value normalized by global %mCH levels. **c**) heatMaps of the top four astrocyte-specific hyper-mCH genes identified using a two-sided Wilcoxon rank-sum test with Bonferroni adjustment for multiple testing (*BMI1 W =* 152892, *p*_adj_ *=* 1.50E-43; *CTTNBP2 W =* 202495, *p*_adj_ *=* 2.78E-83; *MAP3K3 W =* 178268, *p*_adj_ *=* 2.34E-61; *POU4F1 W* = 100516, *p*_adj_ *=* 3.45E-28). Color indicates %mCH for each 500bp genome window averaged by cell type. The gene body is plotted below with exons in black and the predicted promoter site in red. **d**) Top: UMAP distribution of the top astrocyte-specific hyper-mCH gene *BMI1*. Bottom: Average %mCH of all 40 astrocyte-specific hyper-mCH genes normalized by global %mCH. **e**) heatMaps of all four oligodendrocyte-specific hyper-mCH genes identified using a two-sided Wilcoxon rank-sum test with Bonferroni adjustment for multiple testing (BROX *W =* 301188, *p*_adj_ *=* 2.36E-44; *PCDHGC3 W =* 317196, *p*_adj_ *=* 1.74E-63; *PCDHGC4 W =* 309074, *p*_adj_ *=* 1.03E-69; *PCDHGC5 W* = 301393, *p*_adj_ *=* 8.83E-72). Color indicates %mCH for each 500bp genome window averaged by cell type. The gene body is plotted below with exons in black and the predicted promoter site in red. **f**) Top: UMAP distribution of the top oligodendrocyte-specific hyper-mCH gene *BROX*. Bottom: Average %mCH of four oligodendrocyte-specific hyper-mCH genes normalized by global %mCH. **g**) RNA transcript abundance in Allen Brain Atlas subtypes^49,50^ of the 40 genes with astrocyte-specific hyper-mCH (left) and four genes with oligodendrocyte-specific hyper-mCH (right) divided by broad cell class. Values are normalized trimmed mean counts per million averaged by subtype. Column height and error bars indicate group mean ± SE. Values are shown as ln(x + 1) for clarity.

However, in mature glia sharing the same lineage as neurons, Amethyst revealed a novel mCH signature: while hyper-mCH genes in the neuron groups tended to be shared results within similar populations, hyper-mCH genes in astrocytes and oligodendrocytes tended to be specific to their respective groups (**Fig. 4a-b**). Top results for the astrocyte group include *BMI1* (*p*_adj_ = 1.50E-43), a chromatin remodeler with widespread roles in neurodevelopment; *CTTNBP2* (*p*_adj_ = 2.78E-83), a neuron-specific protein involved with dendritic spine formation and implicated in autism spectrum disorder;^51^ and neural transcription factor *POU4F1* (*p*_adj_ = 3.45E-28; **Fig 4c-d**). The oligodendrocyte group only had 4 results – *BROX* (*p*_adj_ = 2.36E-44), a nuclear envelope-associated factor involved in mitotic membrane reassembly; and 3 adjacent clustered protocadherins with reported roles in regulating cortical interneuron programmed cell death (**Fig. 4e-f**).^52^ Visualization of hyper-mCH across these protocadherins shows identical patterns with short trimodal peaks around the beginning, middle, and end of the gene (**Fig. 4e**), suggesting more nuance to mCH deposition than previously appreciated.

While prior research has shown that gene body mCH is anti-correlated with gene expression in neurons,^39^ it is not fully established whether hyper-mCH in glia follows similar principles. To investigate this, we looked at expression of genes specifically hypermethylated in astrocytes or glia across different populations in publicly available Allen Brain Atlas data.^49,50^ For both astrocytes and oligodendrocytes, genes with high mCH accumulation were strongly suppressed in their respective classes, but robustly expressed in neurons (**Fig. 4g**). Genes with the most mCH accumulation tended to be highest expressed in other groups – for example, top hyper-mCH astrocyte gene *CTTNBP2* was barely expressed in astrocytes (mean±SE trimmed normalized expression = 0.008±0.008 CPM), yet extremely abundant at the RNA level in every other group (e.g., 7.18±0.084 CPM in excitatory neuron subtypes). These findings suggest that mCH plays a biologically relevant role in glia as well as neurons and performs a similarly repressive function.

### Atlas-scale data demonstrates glia more often utilize the mCCG trinucleotide context compared to neurons

While other packages for single-cell methylation sequencing exist,^23,24,53–59^ the majority fail when processing datasets of even a few thousand cells. This is a significant obstacle as new protocols such as sciMET^19–22^ easily produce over a hundred thousand cells in a single experiment. To test the efficacy of Amethyst at processing atlas-scale datasets, we analyzed data from Brodmann Area 46 across four human individuals and two sequencing platforms produced using sciMETv3.^22^ Amethyst was capable of processing mCG and mCH data from all 147,901 cells that passed filtering, cleanly separating out distinct neural populations (**Fig. 5a-b**) without sequencing platform or sample origin bias (**Fig. 5c-d**). Benchmarking initial processing steps with successively down-sampled populations revealed a linear relationship between processing time and number of cells (**Fig. 5e**) at nanoseconds per cytosine. While multithreading resulted in substantial memory requirements for large datasets, measures could be taken to reduce computational load if necessary, such as processing chromosomes or samples separately.

**Figure 5.**
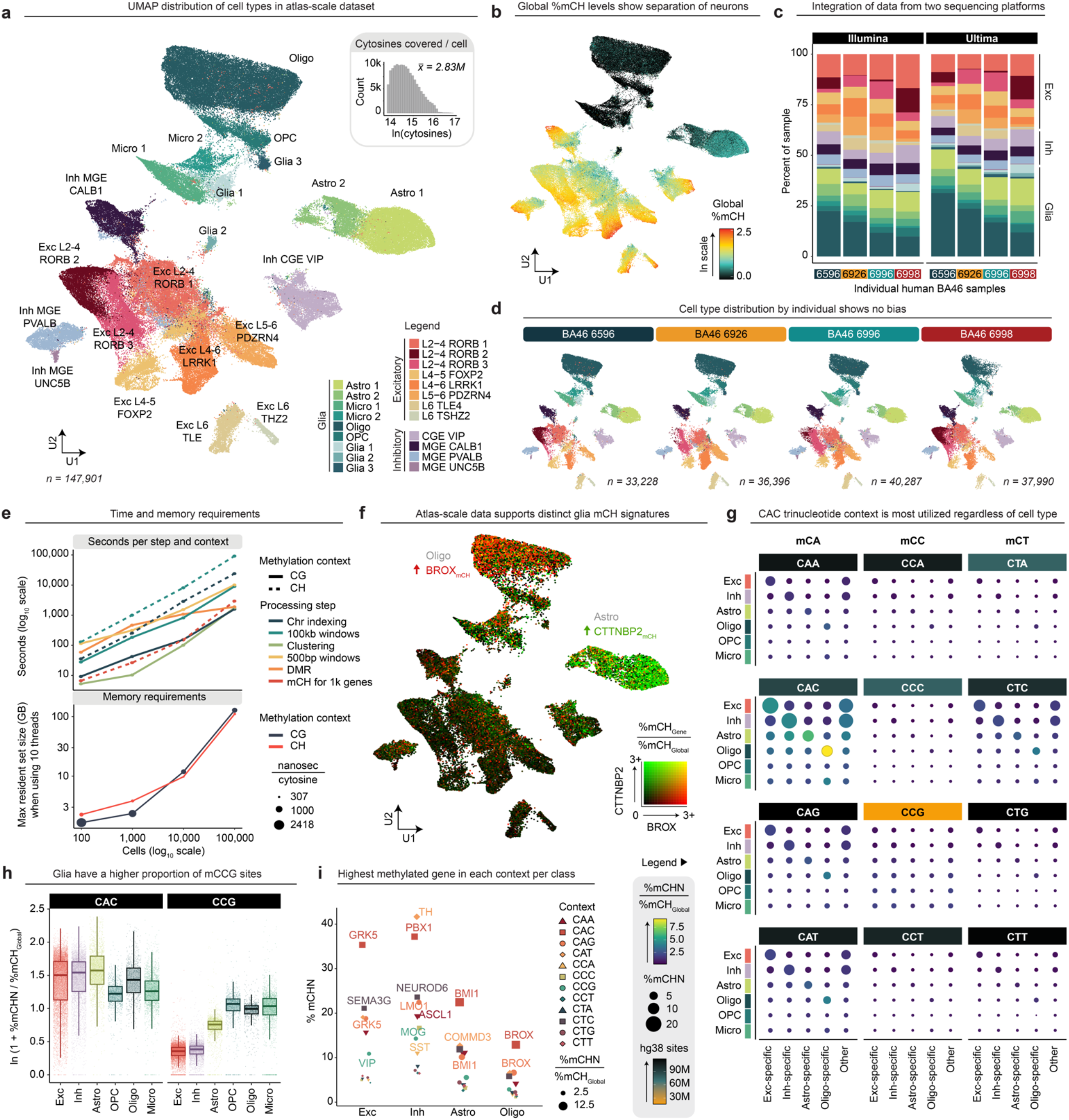
Atlas-scale data demonstrates glia more often utilize the mCCG trinucleotide context compared to neurons. **a**) UMAP plots showing distribution of each cell type from an atlas-scale preparation of 147,901 cells across Brodmann Area 46 (BA46) of four adult human individuals. **b**) dimFeature plot shows the distribution of global %mCH values (on a ln color scale) in UMAP space, revealing clear separation of neurons and glia. **c**) Bar chart showing subtle variation in the cell type composition by individual, but consistent results between Illumina and Ultima sequencing platforms. For a more detailed color legend, see **a. d**) UMAP plots of each sample colored by cell type show no batch effects. **e**) Time and memory requirements for processing successively down-sampled datasets of the top 100k coverage cells in **a** (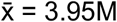 cytosines / cell). Top: Seconds for each process to run at each depth. Color indicates step; line type denotes methylation context. Both axes are on a log_10_ scale. All steps except clustering (IRLBA, Rphenograph, and UMAP steps combined) were run with 10 parallel cores. Bottom: Maximum resident set size in gigabytes for chromosome indexing and window generation (mCH) plus clustering steps (mCG) when using 10 parallel cores. Line color indicates methylation context and dot size indicates the average number of nanoseconds required for processing each cytosine. **f**) UMAP shows oligodendrocytes had highly elevated %mCH levels of *BROX* (in red) normalized by average cell global % mCH (mean±SE: 3.64±0.04 vs. 0.66±0.01; *W* = 267397822, *p* < 2.2E-16 using a two-sided Wilcoxon rank-sum test), while astrocytes had increased normalized %mCH over *CTTNBP2* (in green; mean±SE: 2.11±0.01 vs. 0.62±0.002 mean±SE; *W* = 547193858, *p* < 2.2E-16 using a two-sided Wilcoxon rank-sum test). In the plot, any value >3.0 is capped for clarity. Results are identical across each of four individuals (not shown). **g**) Dot plot representing methylation levels of each of 12 mCH trinucleotide context possibilities for each broad cell class (row) and top genes known to be hypermethylated in that group (column; 10 for excitatory and inhibitory-specific; 40 astrocyte-specific; 4 oligodendrocyte-specific; and nearly 800 other genes previously reported as mCH markers in brain data^11^). Size of the dot represents average % methylation of the corresponding context; color of the dot shows methylation levels normalized by average global %mCH of the group, and color of the context header indicates number of genomic sites identified in hg38. **h**) Methylation status of the CAC and CCG trinucleotide contexts normalized by global %mCH levels for each gene calculated in **g**. Results are divided by broad cell class. Boxplot boundaries represent the first quartile, median, and third quartile values. y axis is ln(x + 1) for clarity. **i**) Gene with the highest %mCH per trinucleotide context and class of the 839 calculated. Shape + color indicates context; size represents %mCH normalized to average global %mCH of the class. Genes with > 45% mCH and non-coding transcripts were excluded from analysis.

Next, we questioned whether this atlas-scale dataset supported our findings of astrocyte and oligodendrocyte-specific hyper-mCH signatures. Data was in consensus with previous conclusions – for example, oligodendrocytes had highly elevated mCH levels of *BROX* (*p*_Wilcoxon_ < 2.2E-16), while astrocytes had increased mCH over *CTTNBP2* (*p*_Wilcoxon_ < 2.2E-16; **Fig. 5f**), despite having much lower global mCH levels (**Fig. 5b**). Since the mCH classification broadly encompasses any cytosine not followed by a guanine, we asked whether this novel glia-specific methylation pattern utilized a different di- or tri-nucleotide context from neurons, which are known to be enriched for mCAC.^10,48,60^ Discrimination of all twelve mCH trinucleotide contexts (**Fig. 5g**) for 800 genes with highly cell type-specific mCH accumulation^11^ revealed that the CAC context was most abundant regardless of cell class or gene specificity. %mCAC in some genes reached an order of magnitude higher than genomic %mCH levels (**Fig. 5h-i**); for example, dopaminergic neuron transcription factor *PBX1* was 37.19±0.96% methylated in the mCAC context of MGE-derived UNC5B inhibitory neurons, relative to the average global mCH levels of 5.66±0.05% (mean±SE; **Fig. 5i**). Other mCA contexts and mCTC were next highest in abundance (**Fig. 5g**). With the exception of mCTC, other CTN and CCN contexts were rarely methylated. Similar distributions were observed across all trinucleotide sequences except for a notable trade-off between mCAC being more abundant in neurons while mCCG was more frequently utilized in glia, despite far fewer genomic positions available in the hg38 reference genome (**Fig. 5g-h**). Altogether, our results suggest the utilization of different mCH trinucleotide contexts is more nuanced than previously appreciated, but similar patterns are present regardless of cell class or gene function.

## Discussion

While other single-cell modalities such as RNA and ATAC have a plethora of robust tools available to aid users in data interpretation,^25–30^ the single-cell methylation field has lagged behind, necessitating significant computational expertise to interact with the data. Here we aim to lower this barrier with Amethyst, the first comprehensive R package for high-throughput single-cell methylation sequencing data analysis. Amethyst provides a flexible workflow suitable for a variety of model systems to fully analyze data from base-level information to interpretation of differentially methylated regions (DMRs).

Using Amethyst, we identified astrocyte and oligodendrocyte-specific hyper-mCH patterns, a non-canonical methylation context that has been underappreciated in glia since an initial report in 2013 by Lister and colleagues.^10^ Here we provide further resolution to their observation of NeuN^-^ cells carrying hyper-mCH in a subset of repressed genes with important roles in neuronal function.^10^ We found that gene-specific hyper-mCH accumulates in mature glia of ectodermal lineage, but not OPCs or microglia. Hyper-mCH genes in astrocytes and oligodendrocytes tend to be exclusive to their respective groups: for example, while *PCDHGC3-5* were previously shown to have hyper mCH in NeuN^-^ cells,^10^ our results show this is entirely due to enrichment in oligodendrocytes. Similar to what has been shown of neuronal mCH, hyper-mCH in glia tends to accumulate across key genes that delineate closely related cell types in a manner anticorrelated with expression.^10,14^ For example, astrocyte hyper-mCH genes *MAP1B, APP*, and *GABBR2* are highly expressed in neurons; while oligodendrocyte hyper-mCH gene *BROX* is highly expressed in both neurons and microglia, but not astrocytes or oligodendrocytes. This observation bolsters the hypothesis that non-CG methylation plays a unique role in fine-tuning gene expression between closely related cell types in the brain^14^ and suggests the principle also applies to glia, in agreement with Lister *et al*.^10^

Upon discrimination of the 12 trinucleotide possibilities contained within the mCH classification, we found that astrocytes and oligodendrocytes have context frequency distributions similar to what has been previously shown in neurons: mCA – particularly mCAC – is most abundant, followed by mCTC, with little methylation observed in other contexts.^48,60,61^ This pattern strongly coincides with the binding capacity for the only known reader of mCH, methyl-CpG binding protein 2 (MECP2),^60,62^ loss of which causes the devastating developmental disorder Rett syndrome.^63,64^ Rett syndrome patients meet initial developmental milestones but experience progressive loss of coordination and cognitive ability. This onset coincides with the early postnatal window mCH starts to be rapidly deposited in maturing neurons,^10^ strongly suggesting the inability to read mCH plays a key role in Rett pathology. While neurons have been the primary focus of Rett research due to the higher relative abundance of MECP2 and mCH, our results support hypotheses of glial dysfunction in Rett syndrome^65–67^ and suggest that – like in neurons – inability to bind mCAC underlies glial pathology.

While mCH trinucleotide context distributions generally mirrored what has previously been shown in neurons,^60^ we found a proportional over-representation of mCCG sites in glia. This is intriguing as MECP2 was shown to lack affinity for mCCN contexts *in vitro*.^60^ One possible explanation is this abundance is passively accumulated by DNMT3A methylating available CpH sites, and the DNA exposed by differing chromatin conformations between neurons and glia have differential proportions of CCG. However, the notably reduced representation of CCG sites in the hg38 genome argues against coincidence. An alternative explanation is there is an unknown reader of mCH with an affinity for mCCG that has not yet been characterized. This hypothesis is supported by previous work comparing MECP2 and DNMT3A knockout mice models,^68^ which concluded MECP2 is only partially responsible for facilitating the functional outcome of DNTM3A-mediated mCH deposition. The prospect of other readers of non-canonical methylation in the human brain is an intriguing avenue for further exploration.

In addition to the biological questions revealed by this analysis, there are many opportunities for expanding Amethyst’s computational toolkit. Areas for future development include: a more automated feature selection process for clustering, improved handling of missing data, trajectory analysis, application to a wider set of species and tissues, incorporation of multiple modalities, and automated cell type annotation. Utilities are regularly being added and will continue to be optimized as the body of single-cell methylation literature expands. This will be catalyzed by Amethyst and other efforts to make single-cell methylation analysis more accessible, enabling a deeper understanding of how methylation guides biological processes across a diversity of contexts.

## Online Methods

### Peripheral blood mononuclear cell (PBMC) and brain tissue sample processing

Example datasets were prepared as described previously.^20^ In brief, human banked PBMCs were purchased from the St. Charles River Labs (PB009C-1/D340161). PBMCs from one 71F individual were taken through the splint ligation version of the sciMETv2 protocol (Nichols et al. 2022,^20^ see Supplementary Note 1) and enriched for regulatory regions using the Twist Human Methylome Panel from Twist Bioscience (105520) as described in Acharya *et al*.^21^ Human middle frontal gyrus samples were obtained from the Oregon Brain Bank and processed using either the sciMETv2 splint ligation or linear amplification protocol.^20^ Following PCR amplification and purification steps, the library was sequenced on an Illumina NextSeq 2000 using a P2 200 cycle kit. Please see Acharya *et al*.^21^ and Nichols *et al*.^20^ for a more detailed description of the methods used to produce the brain and PBMC data covered in Figs. 1-4.

Atlas-scale brain data produced in Fig. 5 is from four human middle frontal gyrus (BA46) NIH Brain Bank samples. Individuals were F38, M48, F27, and M40. Tissue was processed with our advanced iteration of the sciMETv2 splint ligation protocol, sciMETv3, which utilizes a third round of barcoding to exponentially increase throughput.^22^ Detailed sciMETv3 workflow and sample processing methods are available in our corresponding manuscript.^22^

### Initial processing of FASTQ to H5 files

Base-level methylation calls were determined from raw sequence reads using Premethyst, a series of custom scripts (github.com/adeylab/premethyst). Within Premethyst, reads were first demultiplexed using unidex (github.com/adeylab/unidex) by matching to a whitelist of expected barcodes allowing a hamming distance of 2 for each of 3 specified indexes: 2 incorporated during PCR, and a third during tagmentation, which makes up the first 9 bp of read 2. Bases 9–29 of read 2 were then removed, as they contain the transposase mosaic end recognition sequence (ME). Reads were trimmed for adapter sequences using sciMET_trim.pl, followed by alignment using BSBolt (v1.5.0) using the wrapper script sciMET_align_BSBolt.pl. This script runs the aligner with reads 1 and 2 swapped due to the opposite configuration of sciMET adapters compared to traditional bisulfite sequencing adapters, followed with read sorting by name (cell barcode). PCR duplicates were removed for each cell using sciMET_rmdup_pe.pl and then methyl calls for each context (mCG and mCH) were extracted using sciMET_BSBolt_extract.pl. All calls were then wrapped into one .h5 file using sciMET_cellCalls2h5.py. The .h5 file is organized such that the first level contains groups named for each cell barcode, the second level contains groups for each context within each barcode, and the third contains datasets with corresponding call information for each captured cytosine in every cell and context. Other pipelines that produce this output file structure are equally compatible with Amethyst.

### Dimensionality reduction and clustering

#### PBMCs

After base-level calls were wrapped into an h5 file, an empty Amethyst object was generated with *createObject*. Premethyst output files generated during initial processing containing coverage and global methylation metrics were added using helper functions *addCellInfo* and *addAnnot*. To select for high-coverage cells, the metadata slot was filtered to cells containing 10M to 40M cytosines captured using *dplyr::filter* (v1.1.2). The rows corresponding to each chromosome for every cell were cataloged using the *indexChr* function. Following chromosome indexing, methylation score – a normalized measure of deviation from baseline – was calculated over 100kb genomic windows, regulatory regions, and previously-identified DMRs (based on clusters generated using the 100kb window method) using the *makeWindows* function with minimum observation threshold of 2.

For each condition, *runIrlba* was used to calculate truncated singular values (dims = 24 for 100kb windows, 36 for regulatory regions, and 18 for DMRs based on suggested parameters calculated using *dimEstimate*). The function *regressCovBias* was applied to each resulting matrix, which calculates a linear regression for each dimension against the natural log of cytosines covered and returns the residuals. The output was then used for *runCluster(k_phenograph = 30)* to calculate cluster membership and *runUmap(neighbors = 30, dist = 0*.*1, method = “euclidean”)* to compute a uniform manifold approximation and projection. The resulting UMAP coordinates and membership assignments were stored in the metadata slot and explored with plotting functions such as *dimFeature*.

#### Brain (Figs. 1-4)

Similar to initial PBMC processing, an object was initiated with *createObject* and intermediate output files containing coverage and global methylation metrics were added using functions *addCellInfo* and *addAnnot*. High-coverage cells (6M to 100M) were selected in the metadata slot using *dplyr::filter* (v1.1.2). For both the mCG and mCH contexts, the rows corresponding to each chromosome for every cell were cataloged using the *indexChr* function. These indexes were used when calculating percent methylated CpH positions across 100kb genomic sections and CpG score over 100kb windows using the *makeWindows* function. A minimum of 5 observations were required for mCH windows and 2 for mCG.

After calculation of methylation levels over fixed genomic windows, *runIrlba* was used to calculate truncated singular values for both mCH (dims = 23) and mCG (dims = 26) contexts. The function *regressCovBias* was applied to the resulting matrix, followed by *runCluster(k_phenograph = 25)* to calculate cluster membership and *runUmap(neighbors = 25, dist = 0*.*1, method = “euclidean”)* to compute a uniform manifold approximation and projection.

#### Brain (Fig. 5)

Atlas-scale human middle frontal gyrus data was processed by first filtering cells containing fewer than 1M or more than 20M cytosines covered. mCG score and mCH percent were calculated over fixed 100kb genomic windows using the *makeWindows* function. *runIrlba* was then performed to collapse the mCG context to 6 dimensions and mCH to 10. *regressCovBias* was applied to the resulting matrix, followed by Harmony^32^ to reconcile bias introduced by having data from two different sequencing platforms. *runCluster(k_phenograph = 50)* and *runUmap(neighbors = 30, dist = 0*.*0, method = “euclidean”)* functions were used to resolve distinct populations.

### Cell type annotation

Broad cell class for both PBMC and brain data was determined by visually assessing mCG status over canonical marker genes with functions *heatMap* and *histograM*. For brain data, both global %mCH status and %mCH over canonical marker genes were also used to support classification. Further specification was achieved in brain data by comparison to a reference atlas^12^ containing gene body %mCH values for 1502 differentially expressed genes reported in BA10.^12^ Average values for each gene were calculated for the reference data classes and clusters, then *stats::cor* (v4.2.2) was used to calculate pairwise Pearson correlation coefficients. The top annotation label was utilized if the correlation coefficient exceeded 0.8.

### Doublet detection

High-throughput single-cell assays inevitably generate instances of multiple cells sharing the same barcode. The frequency of these instances – termed doublets – can be tuned, but at the cost of final yield. It is therefore optimal to have a computational method for doublet removal in order to facilitate analysis while maintaining throughput. Amethyst includes a tool for doublet detection that adapts principles from approaches developed for other modalities to accommodate sciMET data (**Fig. S1**).

The Amethyst doublet detection tool first involves the creation of a new Amethyst object where the original genomic window matrices are copied over to the new object. From these matrices, a tunable percentage of artificial doublets are generated, where the methylation value is averaged across two randomly selected cells. A comparable number of dropout windows are retained to mitigate effects from increases in coverage. The new matrices are then taken through the dimensionality reduction process with *runIrlba*. This output is utilized to build a prediction model using a random forest approach trained on the artificial doublets. Finally, this model is applied to an object containing the original cells and simulated doublets, producing a doublet score to facilitate filtering after determination of an appropriate threshold. An example in **Fig. S1** using the PBMC dataset demonstrates the efficacy of this approach with much higher doublet scores (mean±SE 0.07± 0.001 vs 0.73±0.008; *p* < 2.2E-16 using a two-sided Wilcoxon rank-sum test) and a higher fail rate (0.44% vs 12.95%; *p* < 2.2E-16 using a two-sided Chi-square test) for artificial doublets.

### Differentially methylated region (DMR) analysis

DMRs for both PBMC and brain data were calculated in the following manner: for every cell type, the sum of member and nonmember c (methylated) and t (unmethylated) observations were calculated in a 1500bp x 500bp sliding window matrix across the genome using the *calcSmoothedWindows* function. Windows with fewer than 10 observations total in member and nonmember groups were removed. For the remaining contingency tables of c and t sums in member and nonmember cells, a variation of a two-sided Fisher’s exact test was used^42^ to calculate a two-tailed p-value approximation using hypergeometric distribution probabilities with a continuity correction with *testDMR*. Adjustment for multiple testing was then performed with *filterDMR* using a Bonferroni correction for each window and group tested and threshold of *p*_adj_ < 0.01 and | logFC | > 2. Since regions of variable methylation can span tens of thousands of base pairs, results for the same group tested and direction of change were collapsed into one DMR using *collapseDMR* with an allowance of up to 2kb between significant regions and minimum length of 1kb.

### Differentially methylated gene analysis

For brain data analysis, %mCH levels across gene bodies were tested in addition to the mCG approach described above. Gene body %mCH for all protein coding genes was calculated using the *makeWindows(nmin = 2)* function. The resulting matrix was used as input for the *findClusterMarkers* function, which utilizes a two-sided Wilcoxon rank-sum test followed by Bonferroni adjustment for multiple testing to compare values from cells within each cluster versus all other cells. This nonparametric test is favored because normality can’t be assumed. Cluster-specific hypermethylations were then identified by filtering results to *p*_adj_ < 0.001 and logFC < -1, followed by using *dplyr (v1*.*1*.*4)* logic to calculate number of representations for each gene in the results table.

### Gene ontology analysis

To determine if genes overlapping DMRs were enriched for any biological processes, we first utilized the *topGO*^46^ *(v2*.*52*.*0)* package to construct an enrichment analysis object using all protein coding genes as background. *topGO::runTest* was performed using the “elim” algorithm with statistic = “fisher”. As the elim algorithm inherently considers hierarchical relationships within the gene ontology structure and eliminates redundant testing, the developers consider the resulting *p-*values corrected.^46^ After running *topGO::GenTable(topNodes = 500, numChar = 50)*, results were filtered for *p*_*elim*_ < 0.01, number of significant genes contributing to the term > 5, and fold change > 2. To generate a treemap in order to further reduce redundancy and more holistically visualize results, we used *rrvgo*^45^ *(v1*.*12*.*2)* to calculate a similarity score matrix between terms and reduce based on semantic relationship with a threshold value of 0.9. This output was then directly plotted with *treemapPlot*. Only parent term text was included for clarity.

## Data Availability

Human peripheral blood mononuclear cell (PBMC) data are available from the NCBI Gene Expression Omnibus (ncbi.nlm.nih.gov/geo/) under accession GSE250282. Human brain sciMETv2 are available from the NCBI Database of Genotypes and Phenotypes (ncbi.nlm.nih.gov/gap/) under accession phs003091.v2.p1. Raw and processed human sciMETv3 data will be available on the NCBI Gene Expression Omnibus (ncbi.nlm.nih.gov/geo/).

## Code Availability

Initial processing of sequencing output to base-level methylation calls can be performed with Premethyst commands, which are available at github.com/adeylab/premethyst. Amethyst is publicly available for installation at github.com/lrylaarsdam/amethyst. Example vignettes are available to demonstrate the major analysis steps in Amethyst for PBMCs - which only have biologically relevant mCG - and frontal cortex, which also has mCH. Analysis code for this manuscript will be made available at github.com/lrylaarsdam/amethyst.

## Acknowledgements

Funding for this work was provided by NIH BRAIN Initiative RF1MH128842 and a Silver Family Foundation Innovator Award to A.C.A. The authors would like to acknowledge Dr. Sonia Acharya for her contributions to generation of the published PBMC data^21^ used in Figures 1-4. We would additionally like to thank other members of the Adey Lab – as well as the Saunders and O’Roak Labs at OHSU - for helpful discussion and feedback. Finally, we would like to thank Dr. Ryan Lister for valuable input on our introduction.

## Author contributions

L.E.R and A.C.A. devised the concept for Amethyst as a unification and expansion of the pre-existing workflow developed by A.C.A. R.V.N. and B.L.O. developed both the sciMETv2-v3 protocols^19,20,22^ and generated all brain and PBMC datasets used in Figures 1-5.^22^ L.E.R. wrote Amethyst code with contributions from S.C. and A.C.A. L.E.R. wrote the manuscript under the guidance of A.C.A. L.E.R. and A.C.A. generated all figures. S.C. and G.G.Y. provided regular feedback on computational aspects of package development. All authors reviewed and approved this manuscript.

